# Evaluating the information content of shallow shotgun metagenomics

**DOI:** 10.1101/320986

**Authors:** Benjamin Hillmann, Gabriel A. Al-Ghalith, Robin R. Shields-Cutler, Qiyun Zhu, Daryl M. Gohl, Kenneth B. Beckman, Rob Knight, Dan Knights

## Abstract

Although microbial communities are associated with many aspects of human, environmental, plant, and animal health, there exists no cost-effective method for precisely characterizing species and genes present in such communities. While deep whole-genome shotgun (WGS) sequencing provides the highest-level of taxonomic and functional resolution, it is often prohibitively expensive for large-scale studies. The prevailing alternative, high-throughput 16S rRNA gene amplicon sequencing (16S), often does not resolve taxonomy past the genus level and provides only moderately accurate predictions of the functional profile; thus, there is currently no widely accepted approach to affordable, high-resolution, taxonomic and functional microbiome analysis. To address this technology gap, we evaluated the information content of shallow shotgun sequencing with as low as 0.5 million sequences per sample as an alternative to 16S sequencing for large human microbiome studies. We describe a library preparation protocol enabling shallow shotgun sequencing at approximately the same per-sample cost as 16S. We analyzed multiple real and simulated biological data sets, including two novel human stool samples with ultra-deep sequencing of 2.5 billion sequences per sample, and found that shallow shotgun recovers accurate species-level taxonomic and functional profiles of the human microbiome. We recognize and discuss some of the inherent limitations of shallow shotgun sequencing, and note that 16S sequencing remains a valuable and important method for taxonomic profiling of novel environments. Although deep WGS remains the gold standard for high-resolution microbiome analysis, we recommend that researchers consider shallow shotgun sequencing as a useful alternative to 16S for large-scale human microbiome research studies.

---

**IMPORTANCE** A common refrain in recent microbiome literature and scientific talks is that the field needs to move away from broad taxonomic surveys using 16S sequencing, and toward more powerful longitudinal studies using shotgun sequencing. However, performing deep shotgun sequencing in large longitudinal studies remains prohibitively expensive for all but the most well-funded research labs and consortia, which leads many researchers to choose 16S sequencing for large studies, followed by deep shotgun sequencing on a subset of targeted samples. Here we show that shallow or moderate-depth shotgun sequencing may be used by researchers to obtain species-level taxonomic and functional data at approximately the same cost as amplicon sequencing. While shallow shotgun sequencing is not intended to replace deep shotgun sequencing for strain-level characterization, we recommend that microbiome scientists consider using shallow shotgun sequencing instead of 16S sequencing for large-scale human microbiome studies.

## INTRODUCTION

Despite the close association of microbial communities with many aspects of human, environmental, plant, and animal health (1–4), it is not currently possible to characterize precisely the species and genes present in a microbial community in a cost-effective manner. The microbial communities of human microbiomes are complex, multivariate and multidimensional, requiring large studies to power novel biomarker discovery and predictive modeling (2,5). Deep whole-genome shotgun metagenomics (WGS) provides highly resolved strain-level taxonomic and functional information, but is generally cost-prohibitive for large-scale studies. Many of the largest microbiome studies to date have been performed via 16S rRNA gene amplicon sequencing (16S), a cost-effective alternative, but 16S typically provides only genus-level taxonomic assignments (6) and rough estimates of the functional repertoire (7,8), limiting the amount of information that can be learned from the data. The purpose of this paper is to evaluate shallow shotgun as an possible cost-effective alternative to 16S sequencing for large-scale biomarker discovery with improved taxonomic resolution and functional accuracy.

A major concern for the use of any microbiome assay is the ability to identify and quantify taxonomic and functional traits from within a complex community. Deep WGS has a number of advantages over 16S for microbiome profiling for these purposes in well-characterized environments; for example, deep WGS of mixed communities, such as the human gut microbiome, has been effective at recovering strain-level polymorphisms and functional traits for abundant strains (9–11). Both shallow and deep shotgun sequencing sequencing are also less subject to amplification bias than 16S because they do not rely on targeted primers to amplify a marker gene (12). However, at the time of writing, WGS sequencing typically costs nearly an order of magnitude more per sample than 16S for library preparation and DNA sequencing.

Although extensive work has been done to characterize how many 16S reads are required for quantifying relevant biological signals, the same has not been done for taxonomic or functional profiling with shotgun metagenomics (Fig. 1A). The depth necessary for sequencing in a particular study depends on the purpose of the study; in many studies, the key goals are to understand which species and functions are present, and to identify biomarkers related to experimental groups or outcomes. To address the important question of how many reads are required to capture species-level taxonomic and functional assignments (e.g. KEGG Orthology groups (13)), we analyzed shotgun sequencing data at various depths. We found that shotgun sequencing can produce similar quality species and functional profiles to deep WGS using few as 0.5 million sequences per sample, as demonstrated on deep whole-genome sequencing (WGS) samples from the Human Microbiome Project (HMP) dataset (14), the HMP mock community (12), a diabetes study (15), simulated human gut microbiomes, and two novel human stool samples on which we performed ultra-deep sequencing of 2.5 billion sequences per sample.

**Figure 1.**
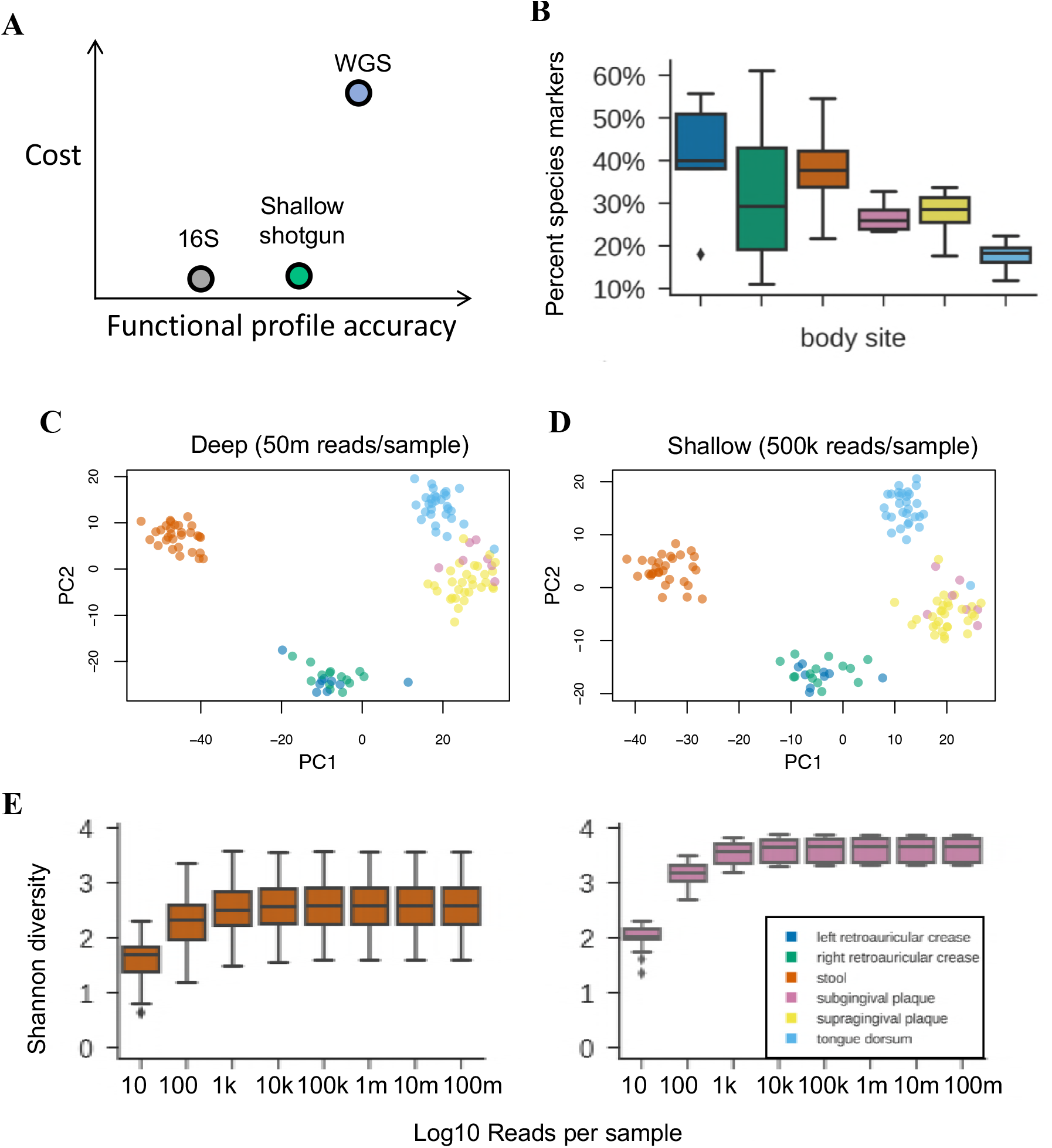
Information content of deep and shallow shotgun sequencing. (A) Shallow shotgun sequencing provides more information than 16S amplicon sequencing, at only slightly higher cost. (B) Percent of raw shotgun DNA sequences that are unique to one bacterial species across different human body habitats (n=7 distinct samples for plaque samples, n=30 distinct samples for other body sites). (C, D) Principal coordinates analysis of Bray-Curtis beta diversity using deep (C) and shallow (D) sequencing (sample sizes as in [B] above). (E) Shannon diversity estimates at varied sequencing depths for human stool (left) and subgingival plaque microbiomes (right; sample sizes as in [B] above). Boxplots show minimum, first quartile, median, second quartile, and maximum, with outliers beyond 1.5 times the interquartile range plotted individually.

## RESULTS

A comparison between deep WGS and shallow shotgun in real and simulated biological data sets demonstrated that shallow shotgun provides nearly the same accuracy at the species and functional level as deep WGS sequencing for known species and genes in five key aspects of microbiome analysis: (1) beta diversity (Fig. 1D-E); (2) alpha diversity (Fig. 1F); (3) species composition (Fig. 2A,C); (4) functional composition (Fig. 2B,D); and (5) clinical biomarker discovery (Fig. 2I).

**Figure 2.**
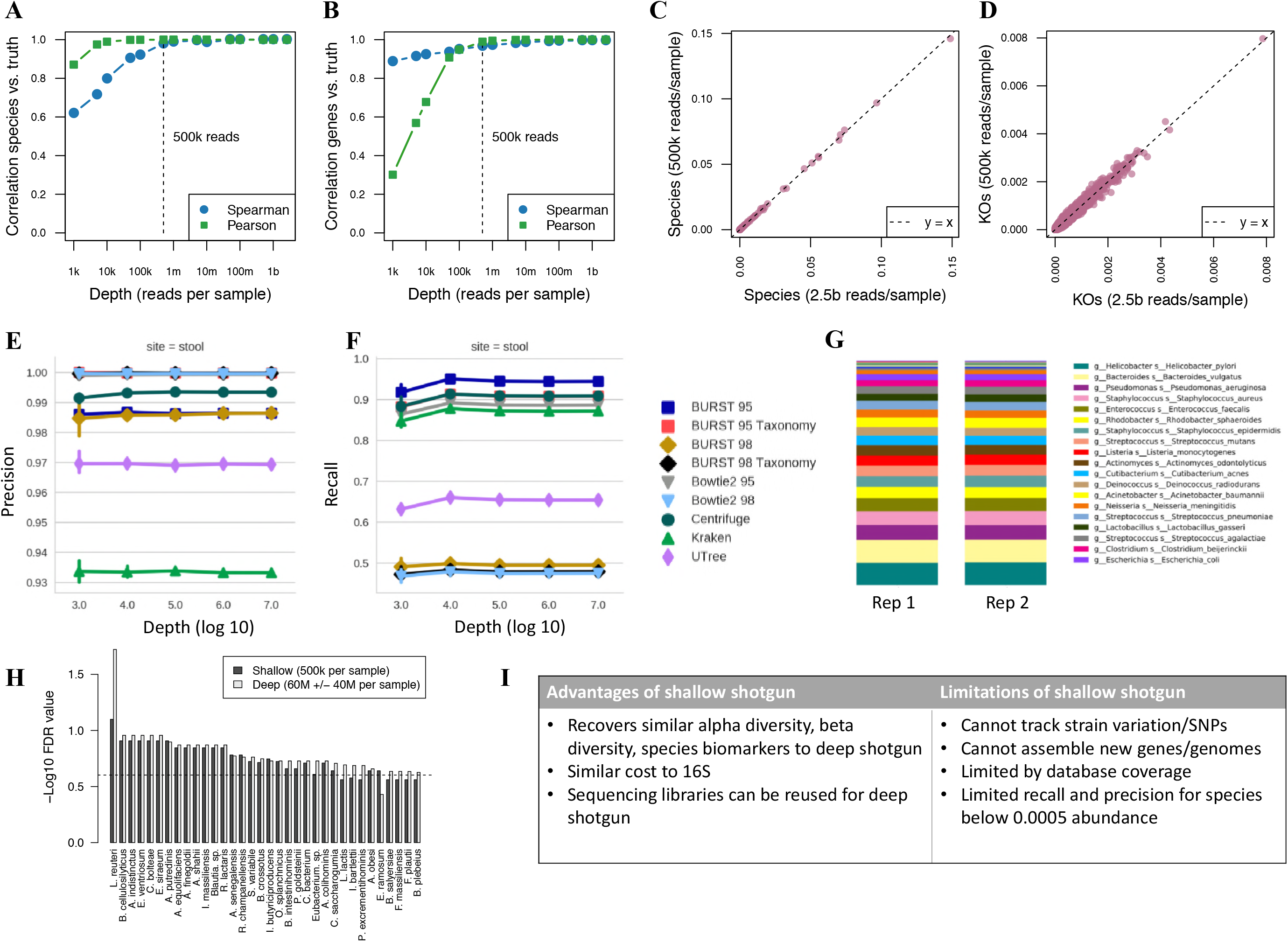
Functional profiling and biomarker discovery using shallow shotgun. (A, B) Correlation with ground-truth species (A) and KEGG Orthology group or KO (B) profile at different sequencing depths, showing that as few as 0.5 million sequences recover nearly the full species and function profiles (ground truth based on 2.5 billion reads per sample; n = 4,394 genes and 694 species at each subsampling level from Subject 1 ultra-deep sequencing sample; comparable results from Subject 2 not shown). (C, D) Scatterplots of species (C) and KOs (D) at 0.5 million versus 2.5 billion reads per sample (same sample size as (A) and (B) above). (E, F) Precision, recall for per-read species binning of different metagenomics analysis tools (“95” and “98” refer to the minimum alignment identity threshold used, n=5 distinct replicates per subsampling depth, error bars show standard deviation). (G) Stacked bar plot of species abundances recovered from HMP mock community shotgun sequencing data. (H) Negative log_10_false-discovery-rate-corrected p-values using Mann-Whitney U tests for species associated with type 2 diabetes (15), compared between deep and shallow shotgun sequencing (n=43 healthy, n=53 type 2 diabetes). (I) Advantages and limitations of shallow shotgun sequencing.

**Alpha and Beta diversity profiling**. We obtained deep WGS data from the Human Microbiome Project (HMP) (14) and sub-sampled the data to simulate shallow shotgun sequencing depth across five body subsites representing skin, oral, and gut habitat microbiomes. Surprisingly little sequencing was needed to discover the same trends found in deep WGS data. To discover these trends, we annotated the deep WGS data using an accelerated version of fully exhaustive gapped Needleman-Wunsch alignment (16,17) for both taxonomic and functional profiles, against all complete, representative bacterial genomes from the reference database RefSeq version number 82 (18). Fully exhaustive alignment allowed us to identify any and all ties for best match for each input sequence according to sequence identity. Species relative abundance profiles were derived by tabulating the number of sequences with at least 80% of the best hits belonging to one species. This is similar to the direct mapping approach used commonly in k-mer-based approaches to shotgun metagenomics taxonomic profiling (19), but with higher sensitivity and recall due to the use of gapped sequence alignment (see simulated data analysis below). Using the fully exhaustive alignment approach, we found that at 20–40% of all sequences could be identified as species markers because they were uniquely present in only one species in the daetabase (Fig. 1B). We then bootstrapped these samples repeatedly down to 10, 100, 1,000, 10,000, 100,000, and 1 million sequences per sample, and reran the analysis to quantify species-level alpha diversity and beta diversity profiles. In all cases, a depth of 0.5 million sequences was more than sufficient to recover the same alpha-and beta-diversity signals as with deep WGS (Fig. 1C-E).

**Species and functional profiles**. In order to compare the performance of shallow shotgun for species and functional profiling with ultra-deep sequencing, we obtained ultra-deep WGS sequencing of 2.5 billion sequences per sample on novel human stool samples from two individuals. At time of writing, to the best of our knowledge, this is the deepest sequencing performed on human stool microbiomes. Using these two novel samples, we measured the species profiles and functional profiles as described above using exhaustive gapped alignment against a full genome database for species and a database of genes annotated with KEGG Orthology groups (KOs) (20). We performed this analysis at the full depth of 2.5 billion sequences, and then subsampled to lower depths. Species profiles at 0.5 million sequences per sample had an average correlation of 0.990 with ultra-deep WGS (Spearman correlation, n=112, p < 2×10^-16^) across the two samples (Fig. 2A,C), and the average KO profile correlation was 0.971 (Spearman correlation, n= 4,394, p < 2×10^-16^) (Fig. 2B,D). For KO annotation we used direct gene observation for all but the lowest abundance genes, where we augmented the direct KO counts with counts of all KOs contained in observed reference strains in a similar manner to Piphillin (8). We weighted the amount of augmentation according to the coefficient of variation of a binomial distribution at the given observed proportion for a given gene with the augmented gene counts contributing at most 10% of the total counts for a given gene. In practice, this approach only affects the most rare genes with fewer than approximately 10 direct observations and offers a slight improvement in accuracy in Spearman correlation with virtually no change to Pearson correlation (see Methods). Our observed 97.1% Spearman correlation of the shallow and deep functional profiles is substantially higher than functional profiles predicted from 16S sequencing, which typically has 80–90% correlation with the directly observed functions (8,7).

Follow the comparison of shallow shotgun to ultra-deep sequencing of real biological samples, we also simulated deep WGS of complex metagenomes from a reference database to evaluate precision and recall of shotgun sequencing at different depths. Individual sequences were drawn at random from full reference genomes of selected species with a simulated 5% rate of sequencing error. Three different mixtures of species were selected from the database to match the average species-level composition of HMP samples from stool, oral, and skin body sites, respectively (see Methods). Precision was defined as the fraction of simulated reads that were correctly assigned to their respective species divided by the total number of reads that mapped to the database. Recall was defined as the fraction of simulated reads that were correctly assigned to their respective species divided by the total number of simulated reads. We found similarly high precision rates of 0.985–0.995 when using exhaustive gapped alignment or Bowtie2 (21) at 95% or 98% alignment identity, or Centrifuge (22); k-mer based methods including Kraken (23) and an in-house method for comparison (24) had lower precision (Fig. 2E). Recall was considerably higher when using 95% identity than 98% identity alignment with BURST or Bowtie2, likely due to the high error rate in the simulated data (Fig. 2F). We also analyzed published shotgun data from the HMP mock community (12), recovering all expected species perfectly as the top 20 taxa, with the exception of *Bacillus cereus* which was recovered at the genus level due to highly overlapping species genomes in the genus *Bacillus* (25) (Fig. 2G).

**Species-level biomarker discovery**. Finally, to assess the ability of shallow shotgun to identify species-level biomarkers in a clinical study, we subsampled deep shotgun sequencing data from a study of healthy individuals and individuals with type 2 diabetes (T2D) (15) to 0.5 million sequences per sample. We identified the species significantly associated with T2D in both the deep data and the shallow data using two-sided Mann-Whitney U tests, and found high concordance between the p-values for species down to 0.0005 relative abundance (average Spearman rho = 0.954 across 10 subsampled replicates, n = 94, p < 2×10^-16^), indicating that 0.5 million sequences per sample enables discovery of species-level biomarkers with comparable power to deep shotgun sequencing down to approximately 0.0005 relative abundance (Fig. 2H). Notably, this classification task contained a range of statistical signals ranging from very strong to marginally significant.

**Comparison to 16S species profiles**. As noted, 16S variable-region amplicon sequences often do not resolve taxa below the genus or family level, although some species can be identified (6). To compare the overall concordance between 16S species profiles and shallow shotgun species profiles for pairs from the same sample, we calculated the Pearson’s correlation R-squared value (coefficient of determination) of the 16S and shallow shotgun species profiles in each pair and found the average R-squared was 0.918. We then permuted the pairing of the 16S and shallow shotgun profiles and repeated the average R-squared calculation to obtain a null distribution, showing that the R-squared between the true pairs of samples was better in all cases than the randomly assigned pairs (Monte Carlo permutation test p < 0.001) (Fig. 3A). This demonstrated high overall concordance between 16S and shallow shotgun species profiles relative to inter-subject differences. To compare the contributions to total relative abundance of observed species between 16S and shallow shotgun profiles, we merged the species-level taxonomic profiles for paired 16S and shallow shotgun analyses and measured the fraction of species attributed to 16S only, shallow shotgun only, or both. We found that there were many species only observed in the shallow shotgun data, with some observed at high levels of abundance (Fig. 3B,C), indicating that the 16S sequencing identified a subset of the dominant taxa at the species level.

**Figure 3.**
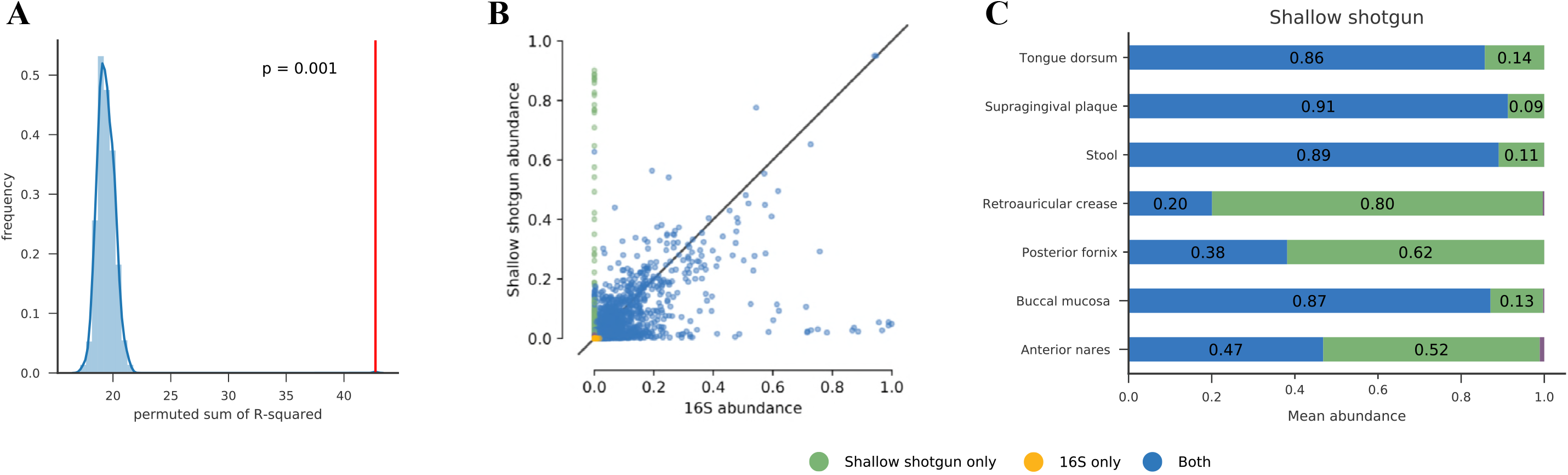
Comparison of 16S and shallow shotgun recovery of species-level taxa. (A) Histogram of average Pearson correlation (R-squared) of species profiles between 16S and shallow shotgun sequencing from the same HMP sample (R2=0.918), compared to the permutation-based null distribution of R-squared values for random pairings (p < 0.001). (B) Scatterplot of relative abundance of species in shallow shotgun sequencing versus 16S sequencing from the same HMP samples; species found only in one data type shown in different color. (C) Fraction of all observed species relative abundance accounted for by species found in 16S only, shallow shotgun only, or both.

## DISCUSSION

In this work we evaluated the information content of shallow shotgun sequencing as a potential alternative to 16S in certain situations. We found that surprisingly few shotgun metagenomic sequences are needed to obtain reliable species and gene group profiles at approximately the same cost as 16S sequencing. We also compared shallow shotgun to deep shotgun on a number of biological data sets including samples from the HMP, a published deep shotgun diabetes study, and simulated and mock communities, and found that we could recover similar trends in alpha and beta diversity, species profiles, and species biomarker discovery down to 0.05% relative abundance with as few as 0.5 million sequences per sample. We then analyzed two human stool samples with new ultra-deep shotgun sequencing data at 2.5 billion reads per sample, the deepest sequencing coverage of any microbiome to our knowledge. We found that shallow sequencing recovers 97–99% correlated species and KEGG (13) Orthology group (KO) profiles when compared to the ultra-deep data.

We did not attempt to perform an exhaustive comparison of different sequence annotation tools as that was outside the scope of our investigation. Instead, we selected several tools representing different approaches to database search for comparison, including exhaustive semi-global gapped alignment (16,17), heuristic gapped alignment using the Burrows-Wheeler transform (21), and k-mer-based search (23,24). Using simulated metagenomic data we found that tools using gapped alignment obtained higher precision and recall than tools using k-mer-based mapping. This result was expected as k-mer mapping requires exact matches of fixed-size k-mers whereas gapped alignment allows insertion of gaps at random to maximize overall sequence identity. In our simulated data, fully exhaustive end-to-end gapped alignment with a minimum threshold of 95% identity using an accelerated version of Needleman-Wunsch (16,17) performed best in terms of recall. Several methods were approximately tied for highest precision. A potential advantage of gapped alignment over k-mer mapping is that current tools report the genomic coordinates of each match, allowing estimation of strain-level coverage, which may be useful for future work into novel algorithms that use strain-level coverage to further improve precision and recall for rare species.

We note a number of important limitations to shallow shotgun sequencing (Fig. 2I). Shallow shotgun sequencing may not be a viable replacement for 16S when characterizing blood or biopsy microbiomes, where there is likely to be more host DNA contamination and relatively low bacterial biomass. Shallow shotgun also relies on whole-genome reference databases, and thus will require expansion of reference genomes to cover novel environments. When analyzing poorly characterized environments, researchers may consider combining 16S sequencing for identification of novel taxonomic groups with shallow shotgun for functional profiling. We have not attempted to compare deep or shallow shotgun sequencing with 16S sequencing in environments with low representation of strains in the reference database, such as marine or soil samples. In these cases it is likely that shallow shotgun will still reveal useful functional profiles due to homology of some observed sequences to known genes and species, but we expect that 16S would provide superior profiling of novel taxa due to lack of available representative genomes covering endemic species, as has been observed for fresh water samples (26).

Shallow shotgun is not meant to be a replacement for deep WGS for strain-level resolution or tracking polymorphisms in strains, and cannot be used for novel gene and genome assembly. For many of the metrics we examined, a depth of 0.5 million sequences per sample was sufficient, but deeper sequencing is warranted for detection of rare species below approximately 0.0005 relative abundance. For this reason, we recommend depths of one or two million per sample for increased sensitivity when possible. In addition, a general concern with any taxonomic annotation is that the boundaries of traditional species taxonomic labels do not necessarily reflect consistent entities at the genomic level when accounting for horizontal gene transfer and inaccurate annotations. These concerns can be alleviated to some extent using deep shotgun sequencing and metagenomic assembly (10), co-abundance clustering (11), or proximity-based assembly (27), although de novo assembly of strains from complex microbiomes remains an active area of research.

We found that shallow sequencing of human stool microbiomes provides high-quality species and functional profiles of human microbiome samples, for little more than the cost of 16S amplicon sequencing when using a miniaturized library preparation protocol (see Methods). We have made available the gene and genome databases that we used together with a convenient Python-based wrapper script that allows users to compare several existing tools for performing both taxonomic and functional annotation (see Methods). Shallow shotgun has a number of important limitations and is not intended to replace deep whole-genome shotgun sequencing for strain-level analysis or novel gene and genome assembly. Nonetheless, shallow shotgun analysis provides considerably more accurate functional profiles and more precise taxonomic resolution than 16S amplicon sequencing for human microbiome studies. Thus, shallow shotgun sequencing is a viable alternative to 16S for researchers performing large-scale human microbiome studies where deep WGS may not be possible.

## Acknowledgements

The authors are grateful to the University of Minnesota Genomics Center for assistance with implementing shallow shotgun DNA sequencing. The research leading to these results has received funding from NIH Grant R01AI121383 (to D.K.), P01DK078669 and in-kind support from Illumina (to R.K.).

## Author contributions

B.H., G.A., and D.K. designed the algorithms. B.H., G.A., D.K., R.S-C., and Q.Z. performed data analysis. D.G. and K.B. designed the wet-lab methods. B.H., D.K., and R.K. wrote the manuscript with contributions from G.A., R.S-C., D.G., and K.B.

## MATERIALS AND METHODS

### Alignment algorithms

Alignment was performed using several existing tools and algorithms, including Bowtie2 (21), Centrifuge (22), Kraken (19), an in-house k-mer based aligner for comparison with Kraken (24) and an accelerated adaptation of Needleman-Wunsch for exhaustive gapped semi-global alignment (16,17).

### Shotgun Species Profiling

After trimming sequences until quality score is above 20, and discarding trimmed sequences shorter than 80 bases or with average quality score of less than 30, query reads were mapped with several different alignment tools against representative and reference genomes from the RefSeq database version 82 (18) using a 95% identity threshold (also compared to 98% for precision and recall evaluation on the simulated data). A read that mapped to a single reference genome is labeled with the NCBI taxonomic annotation. All reads that mapped to multiple reference genomes are labeled as the last common ancestor (LCA) of each label according to the NCBI taxonomy, and only species-level assignments are retained. We use a confidence-adjusted LCA that requires at least 80% of all tied best matches to agree for species annotation. All source code can be found at the github repository: https://github.com/knights-lab/SHOGUN). Additional analysis code used to generate figures and run tests for this manuscript can be found here: https://github.com/knights-lab/analysis_SHOGUN.

### Shotgun Functional Profiling

Functional profiling was obtained using KEGG Orthology (KO) (13) annotations for RefSeq-derived genes (18) from directly observed exhaustive gapped alignments in ultra-deep WGS sequencing. To improve the accuracy of the direct KO profiles for low-abundance genes, the KO profiles were separately predicted from reference genomes and the predicted profiles were used to augment the estimates of low-abundance KOs. Specifically, we identified those query reads with a 100% match to exactly one reference genome, and predicted the entire KO profile of that genome to be present in the sample, similar to a previously published approach (8). This is similar to the PICRUSt algorithm for amplicon sequencing data (7), but without the intermediate steps of clustering short-read amplicons and identifying closely related reference genomes. The final KO profiles reported by SHOGUN are a weighted average of predicted and directly observed KO profiles. The predicted KO counts are weighted between 0.0 and 0.1 by a linear function of the coefficient of variation of the count for a given KO, estimated from the size of the binomial confidence interval for the observed count of a given KO, divided by the count of that KO. The direct KO profiles receive the remainder of the weight, such that the direct KO profiles receive at least 90% of the weight for all genes, and the predicted KO profiles are trusted only for the lowest abundance genes where the expected variance in observed count is high. 16S sequences were aligned to Greengenes version 13_8 (28) at 98% identity with exhaustive gapped alignment (16,17). Where a query sequence aligned equally well to multiple reference sequences, the taxonomic assignment was made using the last common ancestor conserved across at least 80% of the set of references.

### Human Microbiome Project data

We obtained deep WGS data from the Human Microbiome Project (HMP) (14) and sub-sampled the data to simulated shallow shotgun sequencing depth. We annotated the deep WGS data using fully exhaustive gapped alignment for both taxonomy and functional profiles against all complete, representative bacterial genomes from the reference database RefSeq version number 82 (18). We then rarefied these samples repeatedly to 1,000, 10,000, 100,000, and 1 million, and 10 million sequences per sample, and ran the SHOGUN pipeline to quantify species and gene profiles. The list of HMP WGS and corresponding 16S samples used are provided in Supplementary Tables 1 and 2, respectively. The HMP mock community data are from runs SRR2726671 and SRR2726672 from NCBI accession SRX1342165 (12).

### Simulated human metagenomes

The body sites analyzed from the HMP1 project were first grouped according to the broad stool, skin, and oral body sites. We calculated the average relative abundance of all samples within each group. The 100 most abundant species for each group were used for simulating communities. The reads were simulated from a randomly selected strain belonging to each of those most abundant species according to the average proportion of that species in the respective body site group using the tool dwgsim (29). The reads were simulated with default settings for Illumina single-end sequencing machines with a 5% mutation rate where 2% of mutations are indels and a maximum of ten ambiguous bases per query sequence.

### Sequencing library preparation

Shotgun DNA sequencing was performed on the Illumina HiSeq platform. DNA was extracted using the Qiagen DNeasy PowerSoil kit, and was quantified using the Quant-iT PicoGreen dsDNA assay (Thermo Fisher). DNA sequencing libraries were prepared using one-quarter-scale NexteraXT reactions (Illumina). The resulting DNA libraries were denatured with NaOH, diluted to 8 pM in Illumina’s HT1 buffer, spiked with 1% PhiX and a HiSeq 1x100 cycle v3 kit (Illumina) was used to sequence samples. Samples are barcoded and multiplexed on a HiSeq high-output run, with an expected output of at least 0.5 million total sequences per sample. For the ultra-deep shotgun sequencing, 64 separate libraries were prepared as described above but using full Nextera reactions from a homogenized stool sample and were multiplexed on a HiSeq 3000 high-output run, using an entire run per sample.

### Data availability

The data for the ultra-deep WGS sequencing have been deposited in the European Nucleotide Archive with the accession code PRJEB24152.

### Ethics statement

Volunteers contributing stool samples for the two ultra-deep WGS sequencing analyses were recruited as part of research protocol #150275 approved by the University of California San Diego Institutional Review Board. Research was conducted in accordance with the Helsinki Declaration. Informed consent was obtained from all subjects recruited into the study.

